# Predictive Deep Learning Model for Neural Vessel Occlusion

**DOI:** 10.1101/2023.09.27.559850

**Authors:** Aaron Karlsberg, Sandy Kim, Nima Zaghari, Lu Li

## Abstract

Recent advances in medical applications for clot removal have enabled physicians to unblock blood vessel occlusions that occur in extremely narrow regions of the brain along the Horizontal (M1) segment of the middle cerebral artery. FDA approved, clinical trials have proven the significant cerebral recovery that can be achieved for victims of ischemic stroke when they undergo swift surgical intervention for clot removal. That said, carrying out such a delicate operation simultaneously requires extreme surgical expertise, and the ability to quickly identify the site of occlusion from 2D x-ray images of the brain. Fortunately, recent efforts in object detection and classification within the fields of deep learning and computer vision have dramatically improved the predictive power of the computer. Accordingly, the goal of this research project is to develop a deep learning model capable of accurately predicting the site of occlusion from frontal view cerebral angiograms in real time. Our current model utilizes YOLOv3 architecture and identifies the site of occlusion in 94.4% of all cases given a minimum 25% confidence threshold. Furthermore, in 83.97% of cases, the occlusion region is detected with at least 50% average intersection over union between the predicted region and the ground truth region. Finally, distributed over the entire validation set, the average intersection over union between the predicted region and the ground truth region was 74.29%.

## 1 INTRODUCTION

**E**VERY year in the United States, there are more than 795,000 strokes. Ischemic strokes account for 87% of all strokes and arises when a blood vessel becomes occluded by the buildup of fatty deposits (plaques) which inhibit blood flow to the brain [1]. As a result, brain cells begin to die without the necessary oxygen and nutrients supplied by the blood to sustain them.

Although stroke is the fifth-leading cause of death in the United States and causes more serious long-term damage than any other disease [1], fortunately, researchers and medical experts have found that immediate surgical intervention to remove these clot formations can dramatically increase the probability of brain cell recovery in the affected areas.

Conventional methods for demonstrating the location of cerebral vessel occlusions include digital subtraction angiography (DSA) and computed tomography angiography (CTA).

CTA produces three-dimensional X-ray images of cerebral vasculature by encompassing 360 degrees of rotation around the patient’s head and neck during the procedure. Using these three-dimensional images in conjunction with data extracted from blood pressure sensors surrounding the sites of occlusion, researchers have deployed machine learning models to calculate the expected fractional flow rates (FFR) along bifurcations in the blood vessels. This model detects sites of occlusion by highlighting the abnormalities in FFR correlated with particular vascular features [2]. Although CTA is non-invasive and provides enhanced dimensional information, this imaging technique is often costly and time-restrictive, as surgeons are not able to view the imaging in real time while performing surgery on patients.

In time-sensitive scenarios usually following postictal events, DSA is the only method that allows surgeons to navigate their patients vasculature reliably and in real time for immediate clot removal. In this case, DSA is the goldstandard, two-dimensional imaging, test of choice to detect abnormalities in blood flow. DSA uses a contrast dye injected directly into the cerebral vasculature through a catheter that is strategically navigated by x-ray guidance. Once injected, x-ray images of the head reveal the cerebral vasculature while DSA provides a series of images at 2-3 frames per second, detecting the abnormalities in blood flow represented by the contrast dye [3].

In recent years, there has been a massive growth of biomedical data, such as medical images, due to the advances of high-throughput technologies that deem effective computational methods for analysis and interpretation necessary. For one, deep learning is a recent and increasingly popular field of machine learning that employs multilayered deep neural networks to make sense of large-scale data such as images, sounds, and text [4].

Deep learning for medical imaging has proven to be a powerful tool in diagnosis and treatment of various diseases. Traditionally, extracting distinct features from medical images required highly skilled physicians with expertise. Indeed, the complexity and ambiguity of medical images and the lack of knowledge for interpretation has hindered machine learning applications to medicine [4]. However, recently, deep learning methods have been quite successful in the computer vision field through various tasks such as object recognition, localization, and segmentation in medical image analysis.

This paper aims to incorporate deep learning methods to detect large vessel occlusion via digital subtraction angiography images, providing medical specialists the ability to detect the sites of vessel occlusions in real time. We use an algorithm to determine where contrast flow is altered through the abnormalities in vasculature as well as localized key features. As subtle changes from patient to patient may not be detected as easily, the model must be well-trained to identify the the slight variations of human vasculature despite the lack of large amounts of annotated data.

To achieve this goal, we apply the You Only Look Once, version 3 (YOLOv3) deep learning object-detection model framework. Prior detection models apply the model to an image at multiple locations and scales; in contrary, YOLOv3 applies a single neural network to the full image that divides the image into regions and predicts bounding boxes weighted by probabilities for each region [5]. Thus, YOLOv3 detects one object per grid cell, which enforces spatial diversity. Ultimately, this use of a single network evaluation to make predictions in addition to its ability to run locally, make YOLOv3 extremely efficient and accessible in identifying the areas of altered or halted contrast flow in patient DSA images, indicating neural vessel occlusions, in real-time.

## 2 METHODS

### 2.1 Data

The data was provided by the Department of Neurology at the University of California, Los Angeles. 102 frontal view digital subtraction angiograms (DSA) were collected from 102 anonymized patients identified with a vessel occlusion in the M1 segment of the middle cerebral artery (one DSA per patient). These digital subtraction angiograms were delivered in Digital Imaging and Communications in Medicine (DICOM), the standardized medium and file format for medical image communication and information management. The size of the DSAs were 1024 × 1024 pixels. In addition, an Excel workbook file containing the location of the vessel occlusion for each of the patient’s DSA in the form of x and y pixel coordinates was provided.

### 2.2 Image Processing

Each digital subtraction angiogram was recorded at a rate of three frames per second and varied in the total number of frames. In order to compare methods of image processing against each other, we divided the data into three data sets. ImageJ was used to convert each frame in every DICOM into a JPEG file format (.jpg).

#### 2.2.1 Multiple Frame

The Multiple Frame data set comprised all of the .jpg frames extracted from each of the frontal-view DICOM images. This yielded a total of 428 .jpg images. Of these, 382 were used for training and 46 were used for testing and validation.

#### 2.2.2 Single Frame

The Single Frame data set included a single frame from each DICOM image. The single frame was chosen and defined by the moment after the injected contrast agent has reached the site of occlusion determined by the associated x and y pixel coordinates. Each of these single frames were selected manually. In total, there were 102 single frame images. Of these, 92 were used for training and 10 were used for testing and validation.

#### 2.2.3 Average Frame

The Average Frame data set was generated from averaging the frames of each set of JPEG frames obtained from an individual DICOM image file. The average was generated by running a python script, average.py that summed the pixel values at each location and divided by the number of images in the set. In total, there were 102 average frame images. Of these, 92 were used for training and 10 were used for testing and validation.

### 2.3 YOLOv3

You Only Look Once, version 3 (YOLOv3) is a state-of-theart, real-time object detection system.

YOLOv3’s system predicts four coordinates for each bounding box by using dimension clusters as anchor boxes. During training, it uses sum of squared error loss and predicts an objectness score for each bounding box using logistic regression. Unlike other models, YOLOv3 only assigns one bounding box prior for each ground truth object. Each box predicts the classes the bounding box may contain by using multilabel classification through independent logistic classifiers, namely, binary cross-entropy loss. This better models the data as opposed to using softmax, which can impose the assumption that each box has exactly one class, which is often not the case [5].

#### 2.3.1 Darknet-53

As seen in Figure 1, YOLOv3 uses a network performing feature extraction that uses successive 3 × 3 and 1 × 1 convolutional layers with shortcut connections, for a total of 53 convolutional layers, coined as Darknet-53. Darknet53 achieves the highest measured floating point operations per second, better utilizing the GPU and making it more efficient and therefore quicker to evaluate, as opposed to other state-of-the-art classifiers, which often can have excessive layers and therefore, aren’t very efficient.

**Fig. 1.**
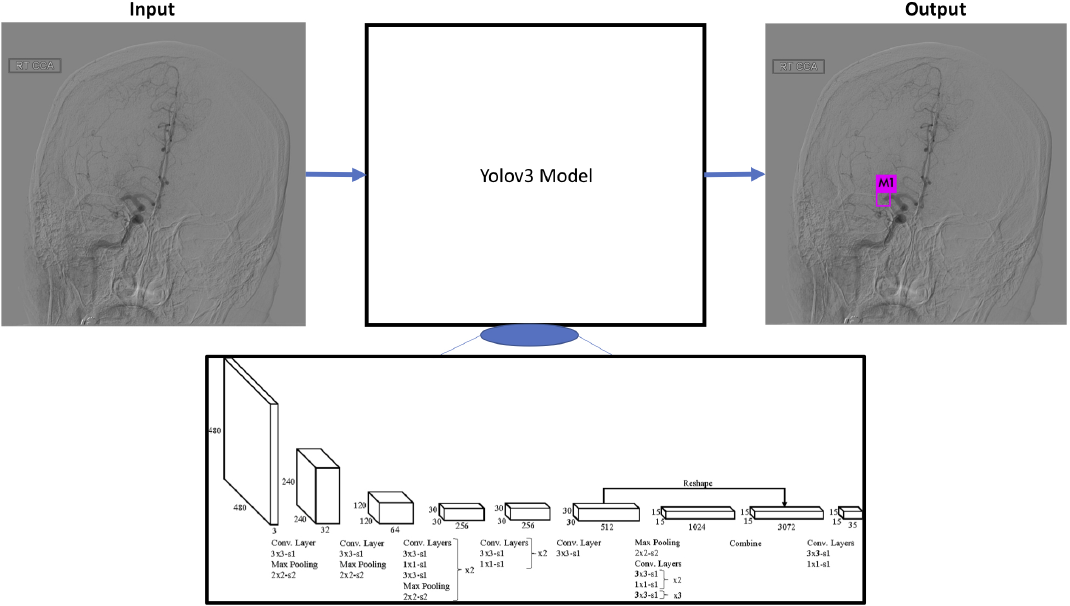
Application of YOLOv3 [6].

YOLOv3 applies the single neural network framework to a full image at test-time, dividing the image into regions and predicting bounding boxes weighted by predicted probabilities for each region, while being informed by global context in the image [5], making it run significantly faster than other detection methods, while its mean average precision remains comparable.

### 2.4 Preparation

#### 2.4.1 Data Annotations

Data labels were created by using Excel to generate a batch (.bat) file and executed on the command line to generate a corresponding text file (.txt) for each and every image.

Each .txt file contained *object class, x, y, width*, and *height*, where *object class* was the index of the object from 0 to classes-1, *x* was the x pixel coordinate of the object, *y* was the y pixel coordinate of the object, *width* was the width of the bounding box, and *height* was the height of the bounding box. In addition, *x, y, width*, and *height* are all float values relative to the width and height of the image.

Data training and validation .txt files were prepared using a python script, process.py.

#### 2.4.2 Configuration Files

Two configuration files were created, a names file (.names) and a data file (.data).

The .names file included the names of the classes, separated by a line break. The .data file included information about the number of classes, the path to the training set, the path to the validation or testing set, the path to the .names file, and the path to where the weights will be saved.

### 2.5 Training

The model was trained using darknet53.conv.74 pretrained convolutional weights with AlexeyAB’s fork of the Darknet neural network framework [7] on an Nvidia GEFORCE GTX 1060 graphics processing unit (GPU).

### 2.6 Evaluation Metrics

To evaluate the performance of the trained model, several metrics were measured. We ran our validation tests on a randomly selected 10% of the data set, while the remaining 90% was used for training.

#### 2.6.1 Average Intersection Over Union

Intersection over union (IoU) is used to measure the accuracy of an object detector on a particular data set. Calculating the IoU requires two sets of data: the ground-truth bounding boxes and the predicted bounding boxes from the model. IoU can then be computed via the ratio

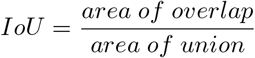

where *area of overlap* is computed between the groundtruth and predicted bounding box and *area of union* is computed as the area encompassed by both the groundtruth and predicted bounding boxes. The mean of the IOU was calculated by accumulating the intersections and unions along all the training instances and then calculating the ratio [8].

#### 2.6.2 Mean Average Precision

Average precision (AP) is another evaluation metric used to measure the accuracy of an object detector that computes the average precision value for recall value over 0 to 1. AP is calculated via the function

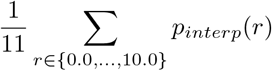

where *p*_*interp*_(*r*) is the maximum precision value for any recall value greater than the recall value for the absolute precision value. AP is then averaged across all categories [9].

#### 2.6.3 Average Loss

Loss is an evaluation metric that measures the performance of a model on its training and validation sets by summing its penalties for bad predictions. Loss is calculated within YOLOv3 via the function

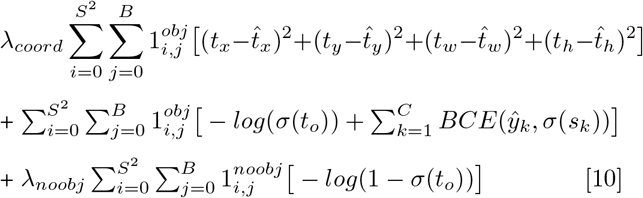

#### 2.6.4 Optimal Weights

Weights were chosen by maximizing both the mean average precision and average intersection over union without having to trade off significantly.

## 3 RESULTS

### 3.1 Multiple Frame

After training on the Multiple Frame data set, the iteration of weights that yielded the highest mean average precision (mAP) and average intersection over union (IOU) based on the validation image set occurred at iteration 10,000. As seen in Figure 2, using this particular iteration of weights our model identifies the site of occlusion in 94.4% of all cases given a minimum 25% confidence threshold. Furthermore, in 83.97% of cases given by the mAP value, the occlusion region is detected with at least 50% IOU between the predicted region and the ground truth region. Finally, distributed over the entire validation set, the average IOU between the predicted region and the ground truth region was 74.29%. The average loss during training at this particular iteration held a value of 0.0196.

**Fig. 2.**
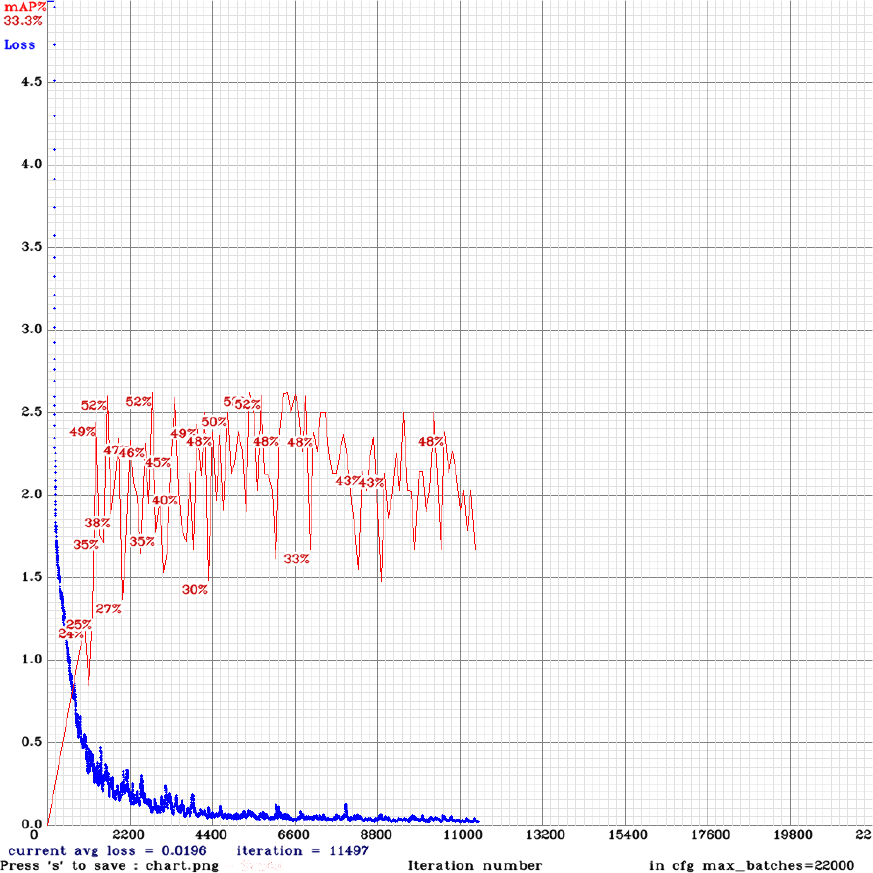
Average Loss and mAP of Multiple Frame Data Set.

### 3.2 Single Frame

After training on the Single Frame data set, the iteration of weights that yielded the highest mean average precision (mAP) and average intersection over union (IOU) based on the validation image set occurred at iteration 8,000. As seen in Figure 3, using this particular iteration of weights our model identifies the site of occlusion in 20.00% of all cases given a minimum 25% confidence threshold. Furthermore, in 10.00% of cases given by the mAP value, the occlusion region is detected with at least 50% IOU between the predicted region and the ground truth region. Finally, distributed over the entire validation set, the average IOU between the predicted region and the ground truth region was 63.02%. The average loss during training at this particular iteration held a value of 0.0199.

**Fig. 3.**
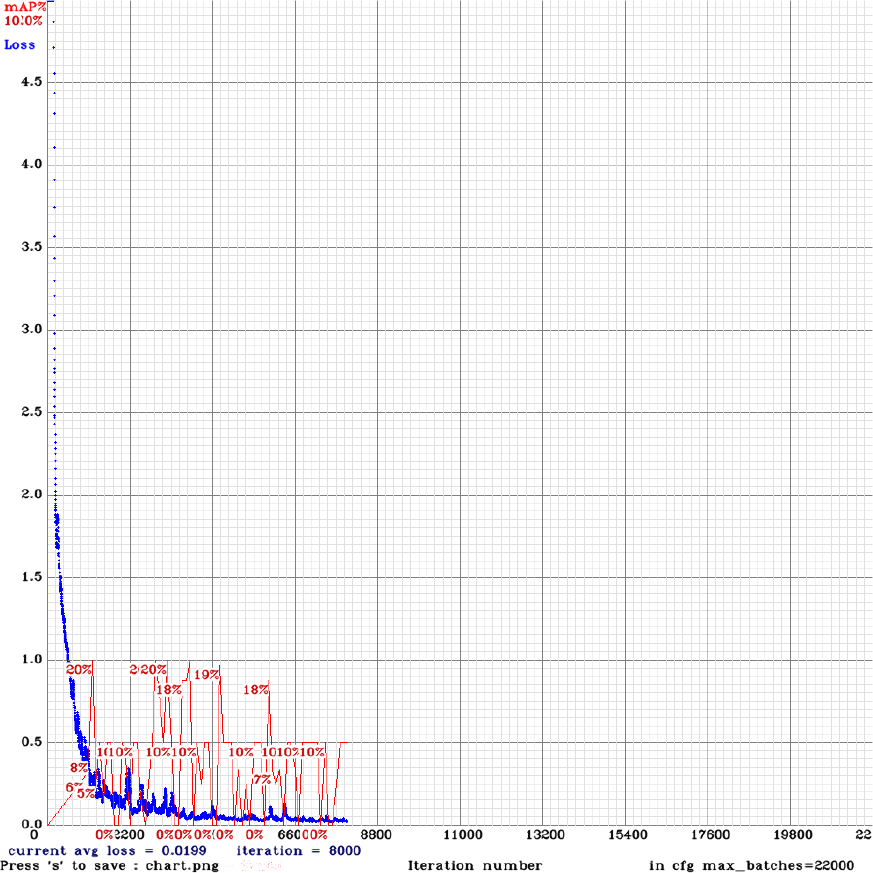
Average Loss and mAP of Single Frame Data Set.

### 3.3 Average Frame

After training on the Average Frame data set, the iteration of weights that yielded the highest mean average precision (mAP) and average intersection over union (IOU) based on the validation image set occurred at iteration 11,000. As seen in Figure 4, using this particular iteration of weights our model identifies the site of occlusion in 50% of all cases given a minimum 25% confidence threshold. Furthermore, in 17.50% of cases given by the mAP value, the occlusion region is detected with at least 50% IOU between the predicted region and the ground truth region. Finally, distributed over the entire validation set, the average IOU between the predicted region and the ground truth region was 63.71%. The average loss during training at this particular iteration held a value of 0.0166.

**Fig. 4.**
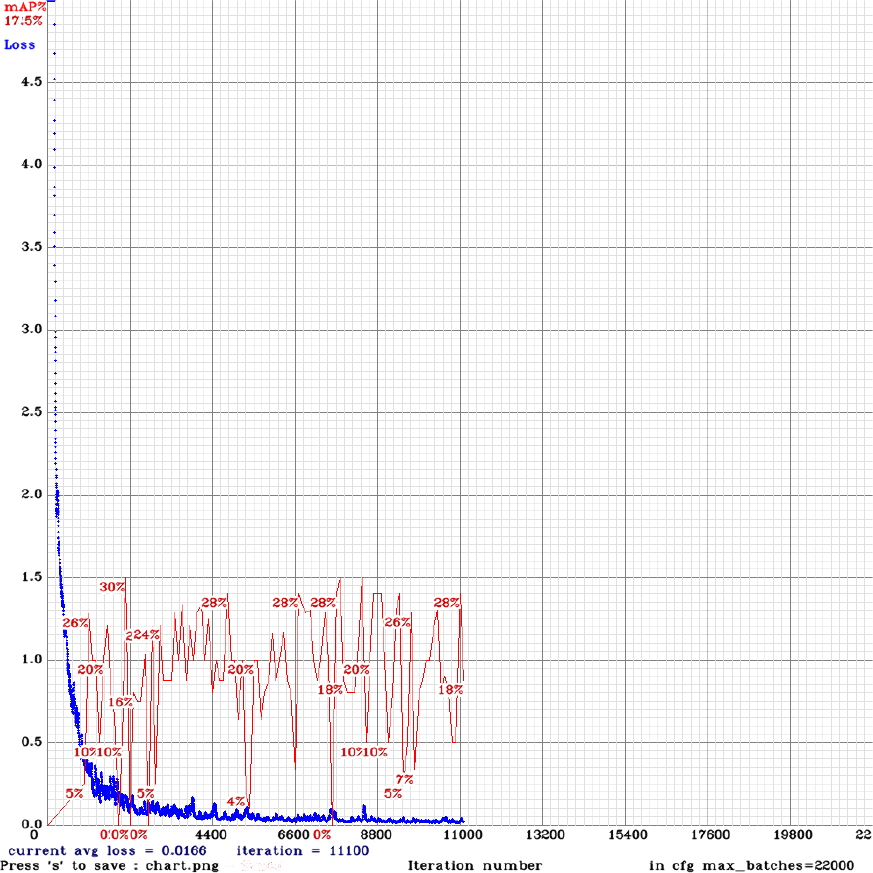
Average Loss and mAP of Average Frame Data Set.

## 4 DISCUSSION

While the results were successful in identifying the location of the occlusion, there still remain many ways to improve the accuracy.

Since our data sets were relatively small, especially Single Frame and Average Frame, adding augmented pictures to the data set would have increased the amount of images to train on and could have potentially provided a higher mean average precision and intersection over union value. Augmenting the images by rotation could give the model a wider array of different positions to detect the image. For example, rotating the image between a range of -30 to 30 degrees would make sense as there can be a slight variation in the orientation of images produced and has been done successfully in similar studies [11]. However, it is probably not necessary to expand outside of that range as images would likely not be flipped completely upside down. Similarly, changing the pixel intensity would be another augmentation that would supplement the data set size. By adjusting the pixel intensity, the model would be better adapted to handle cases where the intensities vary vastly in an image.

In addition, it may be worthwhile to looking at other variables. Parametrizing the vascular structures could enhance the predictive accuracy of the model. For example, we can quantify the blood flow, which would allow us to further label our training data and predict occlusion sites based on both labeled coordinates as well as points of minimal blood flow, such as in previous studies [2].

While this paper focused on developing a model for frontal view digital subtraction angiography images, our overarching goal is to develop another model for detecting occlusions in the lateral view as well. By running these two models simultaneously in the operating room, the surgeon will be able to focus all of their attention on their mechanical task at hand, and rely on our deep learning algorithms for clot detection.

## 5 CONCLUSION

After comparing the performance metrics between the three models, it is clear that the model trained on the Multiple Frame data set performed the best. However, while it is obvious from the results obtained from the Multiple Frame data set that the YOLOv3 deep learning framework is indeed adept at detecting the site of occlusion in frontal view images, we cannot confidently come to a conclusion which methodology of image pre-processing would yield the best predictive model.

The underlying reason for the poor performances of both the Single Frame and Average Frame data sets may be a function of an insufficient quantity training data as opposed to poor quality data. For instance, the Multiple Frame data set contained 4.2x more images than the amount available for training within the Average Frame and Single Frame data sets. It is likely that with enough data, these poor-performing models would substantially improve and perhaps may even exceed the performance of the Multiple Frame model.

However, we were able to generate some insight in comparing the Single Frame versus Average Frame data sets in isolation given that they both used the same amount of training data. From our results it was evident that the Average Frame model performed significantly better than the Single Frame model with a 5-fold improvement in the number of times an occlusion site was successfully detected. This is a sensible outcome given that the Average Frame data set emphasizes all the vasculature along which the contrast agent travels during the surgical operation. As a result, areas that are not emphasized will be associated with occlusion sites. The Single Frame approach likely performed worse for two reasons. First, because it only emphasizes the vasculature surrounding the site of occlusion, rather than a complete picture of the vasculature. Second, because selecting the best frame requires extensive, tedious and manual curation, is slightly subjective, and is therefore prone to inconsistent selection.

Thus, it seems that the best approach moving forward for future studies is to expand training data sets to ensure that differences in performance are not due to data limitations. Future studies may also extend the study by look at training models on different frameworks such as SSD and RNN.

## REFERENCES

[1] Stroke Information, Centers for Disease Control and Prevention. [Online]. Available: https://www.cdc.gov/stroke/. [Accessed: 28-Apr-2019].

[2] Coenen, A., Kim, Y., Kruk, M., Tesche, C., De Geer, J., Kurata, A., Lubbers, M., Daemen, J., Itu, L., Rapaka, S., Sharma, P., Schwemmer, C., Persson, A., Schoepf, U., Kepka, C., Hyun Yang, D. and Nieman, K. (2018). Diagnostic Accuracy of a Machine-Learning Approach to Coronary Computed Tomographic Angiog-raphyBased Fractional Flow Reserve. Circulation: Cardiovascular Imaging, 11(6).

[3] H. A. Reinhold and M. P. West, Nervous System, Acute Care Handbook for Physical Therapists, pp. 123160, 2014.

[4] C. Cao, F. Liu, H. Tan, D. Song, W. Shu, W. Li, Y. Zhou, X. Bo, and Z. Xie, Deep Learning and Its Applications in Biomedicine, Genomics, Proteomics & Bioinformatics, vol. 16, no. 1, pp. 1732, 2018.

[5] Redmon, J. and Farhadi, A. (2018). YOLOv3: An Incremental Improvement. [Online] Seattle, Washington, United States. Available at: https://pjreddie.com/media/files/papers/YOLOv3.pdf [Accessed 28 Apr. 2019].

[6] Aker, C. and Kalkan, S. (2017). Using deep networks for drone detection. [online] Semanticscholar.org. Available at: https://www.semanticscholar.org/paper/Using-deep-networks-for-drone-detection-Aker-Kalkan/64fcde870588026b026b16875fdad67df77d10f9/figure/2 [Accessed 6 Jun. 2019].

[7] GitHub. (2019). AlexeyAB/darknet. [online] Available at: https://github.com/AlexeyAB/darknet [Accessed 6 Jun. 2019].

[8] Rosebrock, A. (2016). Intersection over Union (IoU) for object detection - PyImageSearch. [Online] PyImageSearch. Available at: https://www.pyimagesearch.com/2016/11/07/intersection-over-union-iou-for-object-detection/ [Accessed 6 Jun. 2019].

[9] Hu, J. (2018). mAP (mean Average Precision) for Object Detection. [Online] Medium. Available at: https://medium.com/@jonathanhui/map-mean-average-precision-for-object-detection-45c121a31173 [Accessed 6 Jun. 2019].

[10] Redmon, J., Divvala, D., Girschick, R., and Farhadi, A. (2016). You Only Look Once: Unified, Real-Time Object Detection. [Online] Seattle, Washington, United States. Available at: https://arxiv.org/pdf/1506.02640.pdf [Accessed 6 Jun. 2019].

[11] Unberath, M. Hajek, J. Geimer, T. Schebesch, F. Amrehn, M. Maier, A. (2017). Deep Learning-based Inpainting for Virtual DSA.

